# Impaired social behaviour and molecular mediators of associated neural circuits during chronic *Toxoplasma gondii* infection in female mice

**DOI:** 10.1101/491662

**Authors:** Shiraz Tyebji, Simona Seizova, Alexandra L Garnham, Anthony J Hannan, Christopher J Tonkin

## Abstract

*Toxoplasma gondii* (*T. gondii*) is a neurotropic parasite that is associated with various neuropsychiatric disorders. Rodents infected with *T. gondii* display a plethora of behavioural alterations, and *Toxoplasma* infection in humans has been strongly associated with disorders such as schizophrenia, in which impaired social behaviour is an important feature. Elucidating changes at the cellular level relevant to neuropsychiatric conditions can lead to effective therapies. Here, we compare changes in behaviour during an acute and chronic *T. gondii* infection in female mice. Further, we notice that during chronic phase of infection, mice display impaired sociability when exposed to a novel conspecific. Also, we show that *T. gondii* infected mice display impaired short-term social recognition memory. However, object recognition memory remains intact. Using c-Fos as a marker of neuronal activity, we show that infection leads to an impairment in neuronal activation in the medial prefrontal cortex, hippocampus as well as the amygdala when mice are exposed to a social environment and a change in functional connectivity between these regions. We found changes in synaptic proteins that play a role in the process of neuronal activation such as synaptophysin, PSD-95 and changes in downstream substrates of cell activity such as cyclic AMP, phospho-CREB and BDNF. Our results point towards an imbalance in neuronal activity that can lead to a wider range of neuropsychiatric problems upon *T. gondii* infection.

## 1. Introduction

*Toxoplasma gondii* is a pervasive single-celled intracellular parasite, acquired via the consumption of contaminated food and water, or via vertical transmission during pregnancy, and causes mostly self-resolving flu-like symptoms (Flegr et al., 2014). However, this eukaryotic parasite has a tropism for the brain and muscle tissue, where it establishes a chronic infection. Interestingly, it has been demonstrated that latent *Toxoplasma* infection induces an array of behavioural modifications in rodents including anxiety and depression-like behaviours, and impaired learning and memory (Worth et al., 2014). Such changes are thought to represent adaptive manipulation by the parasite to ensure its transmission back to the feline definitive host, indicating the parasite’s ability to alter a very specific domain of host behaviour (Vyas et al., 2007). Inspired by this, studies in humans have found a strong association with incidence of schizophrenia (SZ) and depression, along with bipolar disorder, general anxiety disorder, aggressive behaviour, acute convulsive epilepsy, suicidal behaviour and self-directed violence (Del Grande et al., 2017; Elsheikha et al., 2016; Esshili et al., 2016; Kannan and Pletnikov, 2012; Sutterland et al., 2015). Despite a wide body of work evaluating the behavioural effects of *Toxoplasma* infection in rodents, there is no clear consensus regarding a mechanism by which the parasite induces such behavioural changes.

Many environmental factors are known to affect a person’s mental health (Schmidt, 2007), of which effects of infectious agents are now well studied (Brown and Derkits, 2010; Fung et al., 2017; Klein et al., 2017). The acute effect of neuroinfections are attributed to inflammatory response and are a hallmark of all infections (Aliberti, 2005; Barichello et al., 2015; Chandran et al., 2011; Ronca et al., 2016), but long-term neurological dysfunction is hypothesized to occur due to irreversible changes that are caused either due to the pathogens altering the host or a persistent host-mediated response to the invading pathogen. Apart from inducing behavioural changes in host (Worth et al., 2014), recent rodent studies report that presence of *Toxoplasma* triggers alterations in existing neuronal pathophysiology, either exacerbating the disease condition (Donley et al., 2016; Mahmoudvand et al., 2016; Montacute et al., 2017) or sometimes even ameliorating (Cabral et al., 2017; Jung et al., 2012) or halting disease progression (Möhle et al., 2016). Two possibilities exist as to how *Toxoplasma* could affect the brain; it could modulate neuronal function either via parasite-induced neuroinflammatory response generated by the host that helps to clear the parasite but elicits secondary neuromodulatory signalling in the brain (Yarovinsky, 2014), or secondly, by direct modification of neuronal signalling by parasitic activity in/around infected neurons (Mendez and Koshy, 2017; Parlog et al., 2015). Either way, it is critical to understand mechanisms by which *Toxoplasma* establishes a chronic infection and specific pathways elicited during the latent infection phase that alters neuronal function that leads to behavioural ramifications and potential neuropsychiatric conditions.

In humans, the strongest link is perhaps between *Toxoplasma* and the susceptibility of individuals to development of SZ. SZ, a chronic, debilitating neuropsychiatric condition, affects approximately 1% of the world population (Saha et al., 2005). Studies have shown that SZ patients are more likely to be seropositive for *Toxoplasma* (Torrey et al., 2007). Not only does *Toxoplasma* infection correlate with the appearance of SZ (Ebadi et al., 2014; Fuglewicz et al., 2017; Torrey and Yolken, 1995; Yolken et al., 2009), but people harbouring latent *Toxoplasma* run a higher risk of developing SZ than any SZ-associated gene variant in the genome wide analysis (Purcell et al., 2009). Murine studies have revealed that *Toxoplasma* infection induces several changes in structure and function of neurons, and alterations in cellular signalling via the dopaminergic (Flegr et al., 2003; Prandovszky et al., 2011; Stibbs, 1985), tryptophan-kynurenine (Silva et al., 2002), Jak/STAT (Kim et al., 2007; Rosowski et al., 2014; Rosowski and Saeij, 2012), GABAergic (Brooks et al., 2015) and the vasopressinergic (Hari Dass and Vyas, 2014) pathways, all of which have been implicated in the pathophysiology of SZ (Brisch et al., 2014; de Jonge et al., 2017; Kegel et al., 2014; Rubin et al., 2014; Singh et al., 2009). Nevertheless, molecular links are yet to be made that directly relates *Toxoplasma* induced brain impairment to SZ-associated behaviours.

Deficits in social behavioural or ‘‘social cognition’’ have been accepted as one of the major symptoms in many neuropsychiatric disorders such as SZ, depression, anxiety, and obsessive compulsive disorder (Derntl and Habel, 2011; Kennedy and Adolphs, 2012). However, social impairments do not necessarily occur in isolation but sometimes have a comorbidity with other mental health disorders such as anxiety (Allsop et al., 2014). In patients with anxiety disorders, social function is significantly affected and is an important symptom when comparing with non-anxious subjects (Kessler et al., 1999; Kroenke et al., 2007). Interestingly, mice infected with *Toxoplasma*, irrespective of strain of parasite, display changes in anxiety (Afonso et al., 2012; Gatkowska et al., 2012; Gonzalez et al., 2007; Hay et al., 1984; Kannan et al., 2010; Machado et al., 2016; Skallová et al., 2006) and therefore, could also display altered social behaviour. This could be a possible link between SZ and *Toxoplasma* infection. Previous studies have shown alterations in social transmission of food preference paradigm (Xiao et al., 2012) and social interactions (Gonzalez et al., 2007) in infected murine models. More recently, Torres et al. reported that 60 days post infection, mice lose their ability to recognise a novel conspecific in a social interaction test (Torres et al., 2018). However, these studies fail to indicate any specific impairment in brain function that could lead to such changes.

Specific signalling mechanisms, such as those involving gene expression and novel protein synthesis, can consolidate short-term memory (STM) into long-term memory (LTM) (McGaugh, 2000). Interestingly, many studies have shown that brain regions displaying learning-induced immediate early gene (IEG) expression play critical roles in such processes (Fukushima et al., 2014; Mamiya et al., 2009; Morrow et al., 1999; Santini et al., 2004; Zhang et al., 2011). Learning and memory processes consist of different phases such as acquisition, consolidation and retrieval (Abel and Nguyen, 2008), and these processes engage different parts of the brain. Similarly, social behaviour entails distinct paradigms of animal behaviour such as social approach, interaction and later recognition or discrimination (Gabor et al., 2012; Lai et al., 2005; Thor and Holloway, 1982). Neuronal firing in brain regions active during such processes can be mapped by looking at factors produced in response to this activity. *c-fos*, a transcription factor, is an IEG that is expressed in an activity-dependent manner (Abraham et al., 1993; Lanahan and Worley, 1998; Montag-Sallaz et al., 1999; Sheng et al., 1990; Worley et al., 1993) and thus has been used to map brain activity upon social interactions (Avale et al., 2011; Filiano et al., 2016; Tanimizu et al., 2017; Wall, 2012). An important master regulator of this signalling mechanism is the transcription factor cAMP-responsive element-binding protein (CREB), that regulates activity-dependent gene expression (Bourtchuladze et al., 1994; Josselyn et al., 2004; Kida et al., 2002; Pittenger et al., 2002). Expression of c-Fos is also mediated by CREB, and thus neural activity seems to initiate a wave of signalling that can regulate both STM as well as LTM (Flavell and Greenberg, 2008), for example transcription of the brain-derived neurotrophic factor (BDNF) (Dong et al., 2006; Katche and Medina, 2017).

For social behaviour, studies have shown that distinct, but interconnected brain regions play an important role in processes such as social interaction and social approach as well as formation/consolidation of social recognition memory (Allsop et al., 2014; Ko, 2017). Important brain regions involved in such behaviours are the hippocampus (Alexander et al., 2016; Hitti and Siegelbaum, 2014; Kogan et al., 2000; Leroy et al., 2017; Lin et al., 2017; Montagrin et al., 2017; Raam et al., 2017; Rubin et al., 2014; Suzuki et al., 2011), the amygdala (Felix-Ortiz and Tye, 2014; Garrido Zinn et al., 2016) and the medial prefrontal cortex (mPFC) (Avale et al., 2011; Franklin et al., 2017; Wall, 2012). Despite one previous report showing impaired social recognition in *Toxoplasma*-infected mice (Torres et al., 2018), origins of such impairment are yet to be understood. In this study, we find that social impairment occurs in mice chronically, but not acutely, infected with *Toxoplasma*. Moreover, infected mice displayed impaired sociability and social memory, but not object recognition memory, indicating that *Toxoplasma* causes changes in specific behavioural domains in the infected host. Using c-Fos as an activity marker, we map neuronal activity in the hippocampus, amygdala and mPFC after social interaction in these mice and show region-specific changes in neuronal activity upon presentation of a novel social stimuli. This is the first report to demonstrate functional impairment in brain regions of *Toxoplasma* infected mice in response to a behavioural task. We also observe changes in proteins mediating synaptic function, particularly in neural circuits involved in learning and memory.

## 2. Methods Mouse model of acute and chronic *Toxoplasma* infection

Six- to eight-week-old female inbred C57BL/6 mice were intraperitoneally injected with tachyzoites of *Toxoplasma* Prugniaud strain (Type II) that were maintained by passage in HFFs. Parasites were harvested using 27-guage needles, pelleted, resuspended in phosphate-buffered saline (PBS) and counted. Mice received either 100 ul PBS (vehicle) or 50,000 tachyzoites resuspended in 100 ul PBS, and were monitored and weighed daily for next 21 days. Mice were housed 4-6 in a box with a 12 hr light-dark cycle, with access to food and water *ad libitum.*

To establish a chronic infection, mice were treated with sulfadiazine sodium in drinking water (100 ug/ml) from day 5 to 10 post-infection. This treatment controls tachyzoite proliferation in the acute stage and helps avoid animal death due to infection.

### Behavioural assays

All behavioural assays were performed in the light phase of the light/dark cycle and only one test was performed in a day. Mice were acclimatised to the procedure room for 1 hr before commencing the tests. Light intensity in the testing room was 100 lux, and the room temperature was kept at 22°C. All test apparatus was cleaned with 70% ethanol and disinfectant F10 between each session and at the end of the last session each day. Where mentioned, the TopScan tracking software (CleverSys, Reston, VA, USA) was used to analyse behavioural videos by an experimenter blinded to the infection status of the animals. Mice that failed to perform a test were excluded from the analysis.

#### Open Field

The open field was a box (40 x 40 cm) made out of white Perspex and was used to analyse anxiety and locomotion in mice. Briefly, mice were placed near the wall and allowed to explore the arena freely for 10 min while being recorded from a top mounted camera. Tracking software was used to analyse location, speed and total distance travelled by the mice. Centre of the open field was defined as 60% of the total area. Light intensity was 50 lux in the centre of the arena.

#### Light-dark box

The light-dark box apparatus consisted of the open field box described earlier with an insert made out of black Perspex that included a window to allow free movement of mice in both directions. At the beginning of the test, mice were place inside the dark area and their movement recorded for 5 min. Video was analysed using the tracking software to determine the number of entries and percentage time spent in the light zone. The light intensity in the light zone was 700 lux.

#### Elevated-zero maze

The maze (San Diego Instruments, San Diego, USA) is a modification of the elevated-plus maze and its design comprises of an elevated (60 cm above the floor) annular platform (10 cm wide), with two opposite enclosed quadrants and two open quadrants, allowing uninterrupted exploration of the maze. This design removes the ambiguity in the interpretation of time spent in the central zone of the more traditional plus maze design (Singh et al., 2007). Mice were placed at the beginning of any one of the closed areas and activity video recorded over a period of 5 min. The analysis software calculated total time spent and entries by the mouse in either the open or closed quadrants. Entry was defined when all four limbs were inside the quadrant.

#### Sucrose preference test

Anhedonia-like behaviour was measured using the sucrose preference test. For this, mice were singly housed and habituated to drinking from two bottles, both containing water. Next day, one of the bottles was switched to 1% sucrose and the weight of the bottles was recorded. The following day, the bottles were weighted again and total water or sucrose consumption was determined. Sucrose preference was calculated as: [sucrose intake/(water intake + sucrose intake)] x 100. The mice were group housed again after the test.

#### Y-maze test for spatial and working memory

The Y-maze apparatus (San Diego Instruments, San Diego, USA) was made of beige-coloured Perspex with three arms (38 cm long) at 120 degrees to each other. The walls of the arms were 12 cm high and 7 cm apart. Two test arms (A and B) contained guillotine doors at their stem to block mouse entry, while the third arm was designated as the home arm (H) with a 10 cm start area.

In the spatial memory task, during the training phase, one of the arms (A or B) was blocked (novel arm) and mice were placed in the start area of the arm H and allowed to explore this arm and the available arm (familiar arm) for 10 min. 1 hr later, to test spatial memory, both arms A and B were made available, and mice were again placed in the start area to explore the maze for 5 min and video recorded. Video was analysed using tracking software to determine time spent by the mice in each arm. Allocation of familiar and novel arms (either A or B) was randomised between each trial. Arm preference was calculated as time spent in each arm x 100/ Total exploration time.

To assess working memory in the Y-maze, mice were placed in the start area of the arm H while the access to other arms (A and B) was open. Sequence of entry into each arm was recorded for 5 min. Entry was defined when all four limbs were inside the arm. One alternation was recorded when the mouse entered a different arm of the maze in each of 3 consecutive arm entries. Percentage spontaneous alternation was then calculated as: number of alternations x 100/total possible alternations.

The spatial memory task was performed at least one week after the working memory task to avoid interference from previous exposure to the apparatus.

#### 3-chambered sociability and social recognition memory task

The 3-chamber test, also known as Crawley’s sociability and preference for social novelty test, is commonly employed to assess the social cognition in mouse models of CNS disorders (Clapcote et al., 2007; Kaidanovich-Beilin et al., 2009; Labrie et al., 2008; Moy et al., 2004). The apparatus (San Diego Instruments, San Diego, USA) consisted of a clear Perspex enclosure with two end chambers (21 x 27 cm) and a central chamber (21 x 12 cm) separated by sliding doors. Test protocol was as described before (Kaidanovich-Beilin et al., 2011). Firstly, test mice were habituated to the empty apparatus for 10 min, and then held in the centre chamber using the sliding doors. Then, an empty wire cup was placed in one of the end chambers and an age and sex-matched control mouse (stranger A) was placed in a wire cup in the other end chamber. To assess the sociability, the sliding doors were opened and mice were allowed to explore for 5 min, and the session was video recorded. Control and test mice were then returned to their home cages. To assess social recognition memory 1 hr later, another age and sex-matched mouse (stranger B) was placed in a wire cup in the end chamber that previously contained an empty cup, and the stranger A was placed as it was earlier. Test mouse was introduced in the central chamber and allowed to explore for 5 min, and the session was video recorded. Time spent by the test mouse sniffing and interacting with either the empty cup (t1), or cups containing strangers’ A (t2) or B (t3) was manually scored. Discrimination/recognition index was calculated as: t2-t1/(t2+t1) or t3-t2/(t3+t2). The chambers used to place the empty cup or cup containing stranger mice was randomised for each test mouse. Before testing, stranger mice A and B were habituated to the wire cup by placing them in the cup for 10 mins, once a day for 3 days.

#### Novel object recognition test

The object recognition test was performed using the same apparatus as the open field, and as described earlier (Tyebji et al., 2015). Briefly, mice were allowed to habituate to the arena for 10 min. Then, in the training phase, two similar objects (A and A’) were placed in the arena (placed diagonally opposite to each other, and equidistant from the centre of the arena) and mice were allowed to explore the objects for 10 min, and the session was video recorded. Mice were then returned to their home cage. To test their object recognition memory after 1 hr (STM), mice were again introduced to the arena for 5 min containing a previously explored familiar object (A or A’) and a novel object (B), and returned to their home cage. To test their LTM, 24 hr later mice were again tested in the arena for 5 min containing previously explored now familiar object (B) and a new object (C). Time spent by the mice sniffing and exploring each object was manually scored. Object preference was calculated as the time exploring each object x 100/ time exploring both objects. Discrimination index was calculated as [(time exploring novel object – time exploring familiar object)/ time exploring both objects].

#### Forced-swim test (FST)

Depression-like behavioural was evaluated using the FST. Mice were individually placed in a 2 ltr glass cylinder filled with water (at 26°C, 15 cm water depth) for 5 min and video recorded. Immobility was defined as remaining motionless, except for certain movements necessary to stay afloat. Latency to first immobility period and total immobility time was manually scored. After the test period, mice were towel dried, placed under a heat lamp for 5 min and then returned to their home cage. FST was the last behavioural assay performed in the battery of tests.

### c-Fos induction after social exposure

Mice were singly housed for 2 days prior to the test and left unmanipulated as previously described (Filiano et al., 2016). Later, infected and PBS injected mice were divided into two groups (total 4), *Social* (n=5) and *Basal* (n=4-8). While the *Basal* group remained in their home cages, *Social* mice were exposed to an age and sex-matched novel mouse kept in a wire cup inside a clear Perspex box (21 x 27 cm) and allowed to interact for 15 min. 2 hr later, all mice were euthanized, transcardially perfused with PBS and their brains removed and stored in 4% paraformaldehyde (PFA) until used for immunohistochemistry.

### Immunostaining of brains sections

PFA fixed brains were paraffin embedded, sliced (5um) using a microtome and mounted on microscope slides before analysis. Staining was performed on the Dako Omnis automated immunostaining platform (Agilent Technologies, CA, USA) and the buffers and reagents mentioned in each step were from EnVision FLEX Systems compatible for use with the Dako Omnis. Briefly, sections underwent dewaxing using the Clearify™ clearing agent and then 2 x wash with distilled water. Target retrieval was performed using target retrieval solution (high pH, c-Fos; low pH, cAMP), washed 2 x with wash buffer, and then incubated with either anti-c-Fos (1:1000, ab190289, Abcam) or anti-cAMP (1:200, ab24851, Abcam) antibody for 1 hr. For c-Fos staining, sections were washed (2x), incubated in peroxidase-blocking reagent (3 min), washed again (2x) and then incubated with biotinylated secondary antibody for 1 hr. After washing 10x, sections were incubated in Streptavidin/HRP reagent for 30 min. Sections were then washed 30 x, incubated with HRP substrate, again washed 30 x, dehydrated and mounted. For cAMP staining, sections were washed 10x, incubated with anti-mouse secondary antibody (1:200, AF594), washed 10 x and then mounted.

### Image analysis

For c-Fos quantification, slides were first scanned using the Pannoramic SCAN II slide scanner and images processed using the CaseCentre slide management application v2.8 (3DHISTECH, Budapest, Hungary). For each brain region, 3-4 regions of interest (ROIs) were acquired at 10x magnification from each animal (1 each from 3-4 individual sections 30 μm apart mounted on the same slide). c-Fos positive nuclei were quantified using Fiji Image J (NIH, Bethesda, USA) and then averaged for each animal (n=4-7). Data is presented as number of c-Fos positive cells counted in each brain region as a percentage of that from animals of the PBS injected *Basal* group.

For cAMP quantification, images were acquired using the Zeiss Axiovert 200M wide field microscope equipped with a AxioCam MRn CCD detector. For each brain region, 3 ROIs were acquired from 3 individual sections from the same animal spaced 30 μm apart which were mounted on the same slide. Staining intensity was quantified using Fiji Image J (NIH, Bethesda, USA) as mean grey value of each ROI and normalized after subtracting background intensity. Values were averaged for each animal (n= 4-7) and data represented as mean florescence intensity as a percentage of that from animals of the PBS injected group.

### Pearson’s *r* correlations

Pearson’s correlations were calculated between brain regions for each experimental group. For all groups except the PBS mice (home cage) group, this was done using the *rcorr* function from the R Hmisc package (Harrell et al., 2018). Asymptotic *P*-values for the significance of this correlation were also calculated by this function. Correlations for the PBS mice (home cage) group were calculated using the base R function *cor*. This was done due to a smaller sample size available in this group. The correlation plots were produced using the R corrplot package (Wei and Simko, 2017).

### Total protein extraction

*Toxoplasma* infected and PBS injected mice at 8 weeks post-infection (wpi) were euthanized by CO_2_. Brains were quickly removed and the hippocampus, amygdala and PFC were dissected out and homogenised by sonication in lysis buffer (RIPA) containing 50mM Tris-HCl (pH 8.0), 150mM NaCl, 1% NP-40, 0.5% Sodium deoxycholate, 0.1% Sodium dodecyl sulphate (SDS), 1mM phenylmethylsulphonyl fluoride (PMSF) and protease inhibitor cocktail (Roche). Samples were centrifuged at 12000 x g at 4°C for 20 min, supernatants collected and protein concentration was determined using the Pierce BCA protein assay kit (Thermo Scientific).

### Western blot analysis

Proteins were denatured in loading buffer containing 50 mM Tris-HCl (pH 6.8), 2% SDS (w/v), 10% glycerol, 0.25 mM EDTA, 2.5% ß-mercaptoethanol and 0.02% bromophenol blue (w/v) by heating at 100°C for 5 min. 30 ug (15 ug for phospho-CREB analysis) protein was resolved in denaturing polyacrylamide gels and then transferred onto nitrocellulose membranes (Whatman® Schleicher&Schuell, Dassel, Germany), washed in wash buffer containing Tris-buffered saline and 0.1% Tween-20 (TBS-T), and blocked at room temperature for 1 hr using TBS-T containing 5% bovine serum albumin and 5% skimmed milk. Membranes were then incubated overnight at 4°C in primary antibodies (1:1000, unless otherwise stated): anti-BDNF (ab108319, Abcam), anti-Synaptophysin (1:500, ab8049, Abcam), anti-PSD-95 (ab18258, Abcam), anti-phospho CREB^ser133^ (05-667, Millipore). Analysis of phospho-CREB levels were performed in membranes blocked by incubation with 5% Phospho-BLOCKER (Cell Biolabs, San Dieogo, CA) in TBS-T. Loading control was performed by re-probing the membranes with anti-ßIII Tubulin (G7171, 1:20000, Promega) during 15 mins at room temperature. After primary antibody incubation, membranes were washed with TBS-T and incubated with appropriate secondary antibody for 1 hr at room temperature (anti-mouse horseradish peroxidase conjugated for synaptophysin and pCREB; anti-rabbit LI-COR IRDye® secondary antibody for BDNF and PSD-95) and the reaction was visualized on a photographic film using the Amersham ECL western blotting liquid (GE Healthcare, UK) or the Odyssey infrared imaging system (LI-COR Biosciences, USA). Densitometric analysis of bots was done using Fiji Image J (NIH, Bethesda, USA).

### Ethics statement

All animal experiments complied with the regulatory standards of and were approved by the Walter and Eliza Hall Institute Animal Ethics Committees under approval number 2014.021.

### Statistical analysis

Data analysis was performed using the GraphPad Prism version 7 (GraphPad Software, La Jolla, USA). Statistical analysis was performed using the Student’s *t*-test or the two-way analysis of variance (ANOVA) followed by Sidaks’s multiple comparison post-hoc test and indicated in the figure legends. A 95% confidence interval was used and values of *P* < 0.05 were considered as statistically significant.

## 3. Results

### 3.1 Behavioural changes in mice during acute *Toxoplasma* infection

Infection with pathogens lead to acute inflammatory responses that are known to affect the mental health status of the host (Klein et al., 2017). Infection with *Toxoplasma* leads to a powerful immune response in order to restrict parasite dissemination and prevent host mortality (Aliberti, 2005; Yarovinsky, 2014). Nevertheless, it is known that inflammatory molecules generated during this period can have affect brain function, leading to changes in host behaviour. To assess the effects of an acute *Toxoplasma* infection on mice, we intraperitoneally injected female C57BL/6 mice with PBS (vehicle) or 50,000 tachyzoites of the type II Pru strain.

At 3 weeks post-infection (wpi), we measured the classic behavioural phenotypes that are associated with *Toxoplasma* infection and also are characteristic behaviour changes observed upon agents that induce inflammatory response in mice (Couch et al., 2016; Cunningham et al., 2009; Tarr et al., 2012; Walker et al., 2013). In a circular open field, when compared to PBS mice, *Toxoplasma* infected mice spent significantly less time (*P* = 0.0015) and made less entries (*P* = 0.0002) in the centre area (Fig. 1A), while in a light dark apparatus, the infected mice spend significantly less time (*P* = 0.0256) and made less entries (*P* = 0.0083) in the light area (Fig. 1B), indicating an increased anxiety phenotype in these mice. But, in the EZM task, we found no change in the time spent in the open arms, or in the entries made into the open arms (Supp. Fig. 1B). Moreover, infected mice covered less distance (*P* = 0.0274) in the open field, while their average speed remained unchanged (Fig. 1C), indicating reduced locomotor activity. Further, we found that infected mice consumed less sucrose water (*P* = 0.0059) (Fig. 1D), showed reduced latency to immobility (*P* = 0.0057) and increased total time immobile (*P* = 0.0487) (Fig. 1E) in the forced-swim test, indicating increased anhedonia and depression-like behaviours at this post-infection stage (3 wpi).

**Figure 1:**
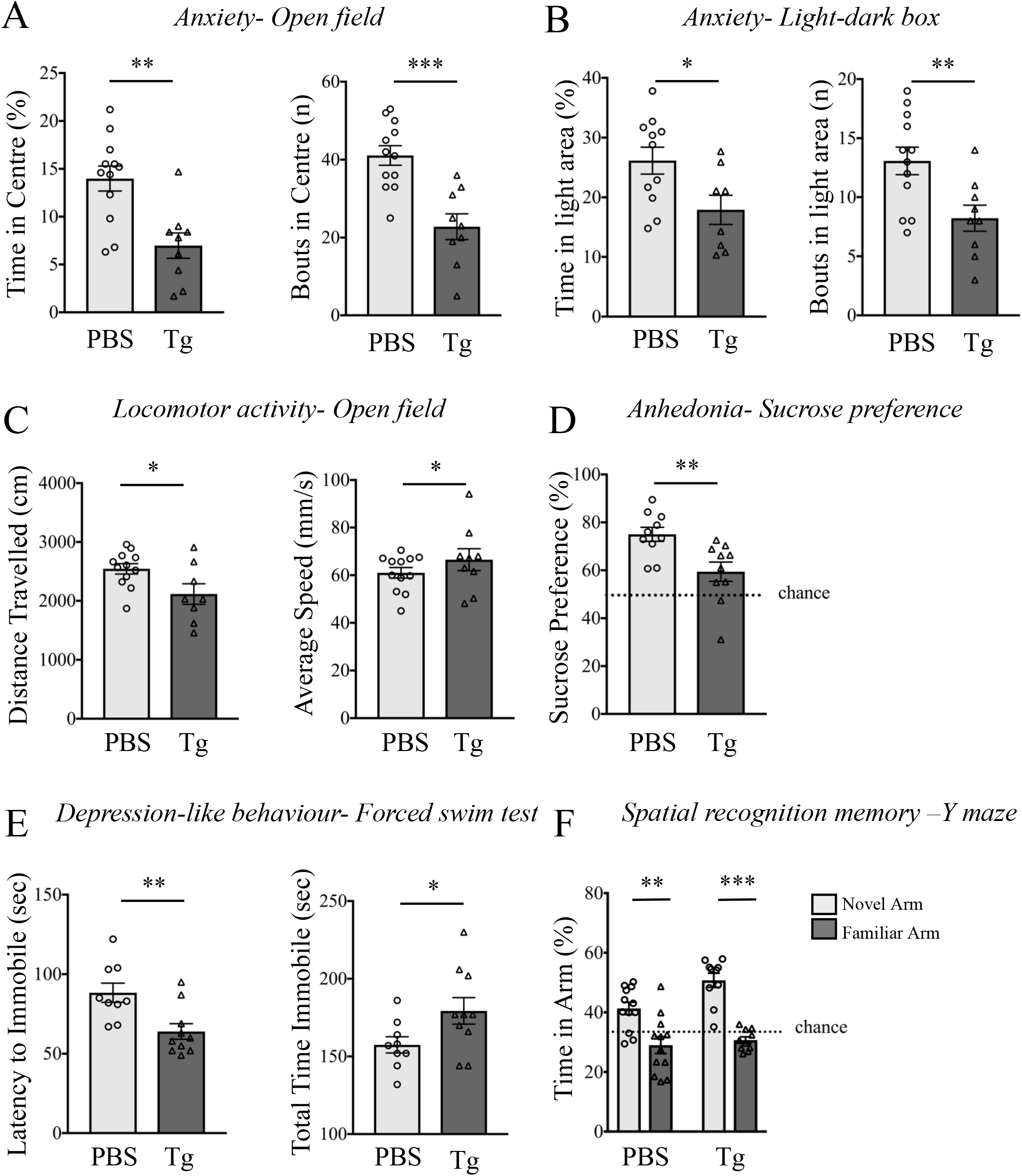
Behavioural changes in mice 3 weeks after *T. gondii* infection. 6 to 8-week-old female C57BL/6 mice were infected with *T. gondii* (Tg) or mock infected (PBS) and behavioural analysis began 3 weeks later. The anxiety phenotype was measured using the open-field test (A) and the light-dark box (B), by calculating the percentage times spent and entries made into the centre area or the light zone, respectively. Locomotor activity was measured as the distance travelled and average speed in the open field (C). Anhedonia was measured by calculating preferential sucrose consumption over water (D). Depression-like behaviour was analysed by measuring the latency to immobility and the total time immobile in the forced-swim test (E). Spatial recognition memory was measured using the Y-maze as time spent in each arm 1 hour after training (F). In all graphs, error bars represent mean ± SEM (n= 9-12). Data were analysed by Student’s *t*-test (A-E) or two-way ANOVA with Sidak’s multiple comparison test as a *post hoc* (F). \**P*<0.05, \*\**P*<0.01 and \*\*\**P*<0.001.

When we measured cognitive changes in mice at this stage of infection, we found no change in the arm preference in the Y-maze test for spatial memory (Fig. 1F) and no change in total alternations in the Y-maze continuous alternation test for working memory (Supp. Fig. 1A) in *Toxoplasma*-infected mice.

To ascertain whether the behavioural changes we observe are due to an acute effect of sulfadiazine treatment provided to the infected mice, uninfected PBS-injected mice underwent a similar sulfadiazine treatment regime and were tested for behavioural changes (Supp. Fig. 3). We only observed an increase in locomotor activity (*P* = 0.0017 and *P* = 0.001 for average speed and total distance covered, respectively) when compared to control mice, suggesting that locomotor changes in *T. gondii* infected mice could be affected by treatment with sulfadiazine.

### 3.2 Behavioural changes in mice during chronic *Toxoplasma* infection

Molecular changes occurring in the brain during an acute inflammatory period may alter the brain neuropathology in a way that can persist over time and therefore lead to changes in brain function long after the infection has been resolved. It is hypothesised that changes in host behaviour observed during a chronic *Toxoplasma* infection could be a cumulative effect of the inflammatory assault that the brain faced during the acute infection phase, combined with the result of subsequent parasitic activity (Parlog et al., 2015). To evaluate whether behavioural changes persisted in our models of infection at chronic stages and whether any further cognitive impairment occurred at this stage, a separate cohort of mice at 8 wpi underwent the same battery of behavioural tests as those which had been performed at 3 wpi.

Infected mice displayed increased anxiety as they spent significantly less time (*P* = 0.0042) and made less entries (*P* = 0.0467) in centre in the open field test (Fig. 2A). Moreover, in the LDB test, infected mice made fewer entries (*P* = 0.042) in the light zone (Fig. 2B). However, we found no significant difference in the time spent in the light zone (Fig. 2B), nor any difference in performance in the EZM test (Supp. Fig. 2B), when compared to PBS mice. Infected mice also showed increased average speed (*P* = 0.0355) in the open field, but no change in the total distance covered (Fig. 2C). We observed that at this stage of infection, mice displayed anhedonia-like behaviour (*P* = 0.0293) (Fig. 2D) but no depression-like behaviour (Fig. 2E) as evaluated by the FST.

**Figure 2:**
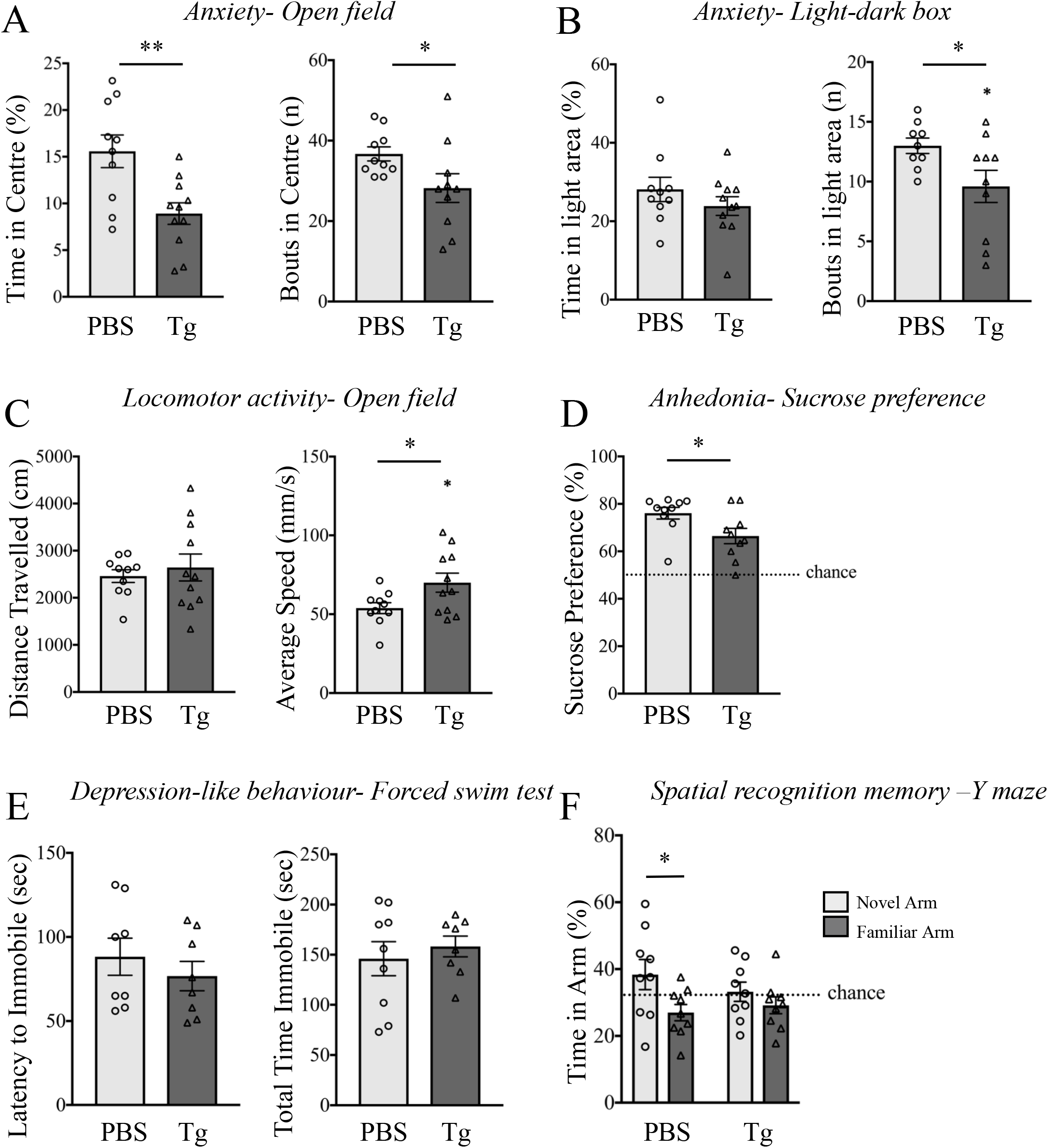
Behavioural changes in mice 8 weeks after *T. gondii* infection. 6 to 8-week-old female C57BL/6 mice were infected with *T. gondii* (Tg) or mock infected (PBS) and behavioural analysis began 3 weeks later. The anxiety phenotype was measured using the open-field test (A) and the light-dark box (B) by calculating the percentage times spent and entries made into the centre area or the light zone, respectively. Locomotor activity was measured as the distance travelled and average speed in the open field (C). Anhedonia was measured by calculating preferential sucrose consumption over water (D). Depression-like behaviour was analysed by measuring the latency to immobility and the total time immobile in the forced swim test (E). Spatial recognition memory was measured using the Y-maze as time spent in each arm 1 hour after training (F). In all graphs, error bars represent mean ± SEM (n= 8-11). Data were analysed by Student’s *t*-test (A-E) or two-way ANOVA with Sidak’s multiple comparison test as a *post hoc* (F). \**P*<0.05 and \*\**P*<0.01.

Interestingly, when we measured spatial memory of mice using the Y-maze, we observed that infected mice displayed no preference for the novel arm (*P* = 0.611) as opposed to PBS mice (*P* = 0.0338) (Fig. 2F). Nevertheless, we failed to obverse any change in their working memory using the Y-maze continuous alternation test (Supp. Fig. 2A).

### 3.3 Chronic *T. gondii* infection impairs social behaviour in mice

Rodents are social animals and interaction and recognition of conspecifics play an important role in maintaining social hierarchy and finding a mate (Berry and Bronson, 1992). Moreover, comorbidity between appearance of anxiety and social impairment in an individual (Allsop et al., 2014) suggests that anxiety phenotype generated due to *T. gondii* infection could also alter the social behaviour in infected mice. Therefore, we measured the sociability in *T. gondii* infected mice using the 3-chambered social interaction test (3CSIT) during acute as well as chronic stages of infection. This assay quantifies the preference of a mouse towards either another mouse or an object, and has been shown to identify social dysfunction in mouse models of disorders that present social deficits (Kennedy and Adolphs, 2012; Silverman et al., 2010). After habituation to the apparatus, when mice were given a choice between an empty cage and a cage containing a novel conspecific at 3 wpi, infected as well as PBS control mice spent significantly more time sniffing and exploring the cage containing a mouse as compared to an empty cage (*P* = 0.0003 and *P* = 0.0153, respectively) (Fig. 3A). To evaluate the strength of sociability, we measured the discrimination index in this test and observed that there was no difference in the discrimination index shown by PBS and infected mice (Fig. 3B). But when we tested mice 8 wpi at the 3CSIT, infected mice did not show any significant preference for the cage containing the mouse (Fig. 3C). Moreover, the infected mice showed a significantly worse discrimination index (*P* = 0.0285) compared to PBS mice (Fig. 3D). Thus, chronic *T. gondii* infection impairs social preference in mice.

**Figure 3:**
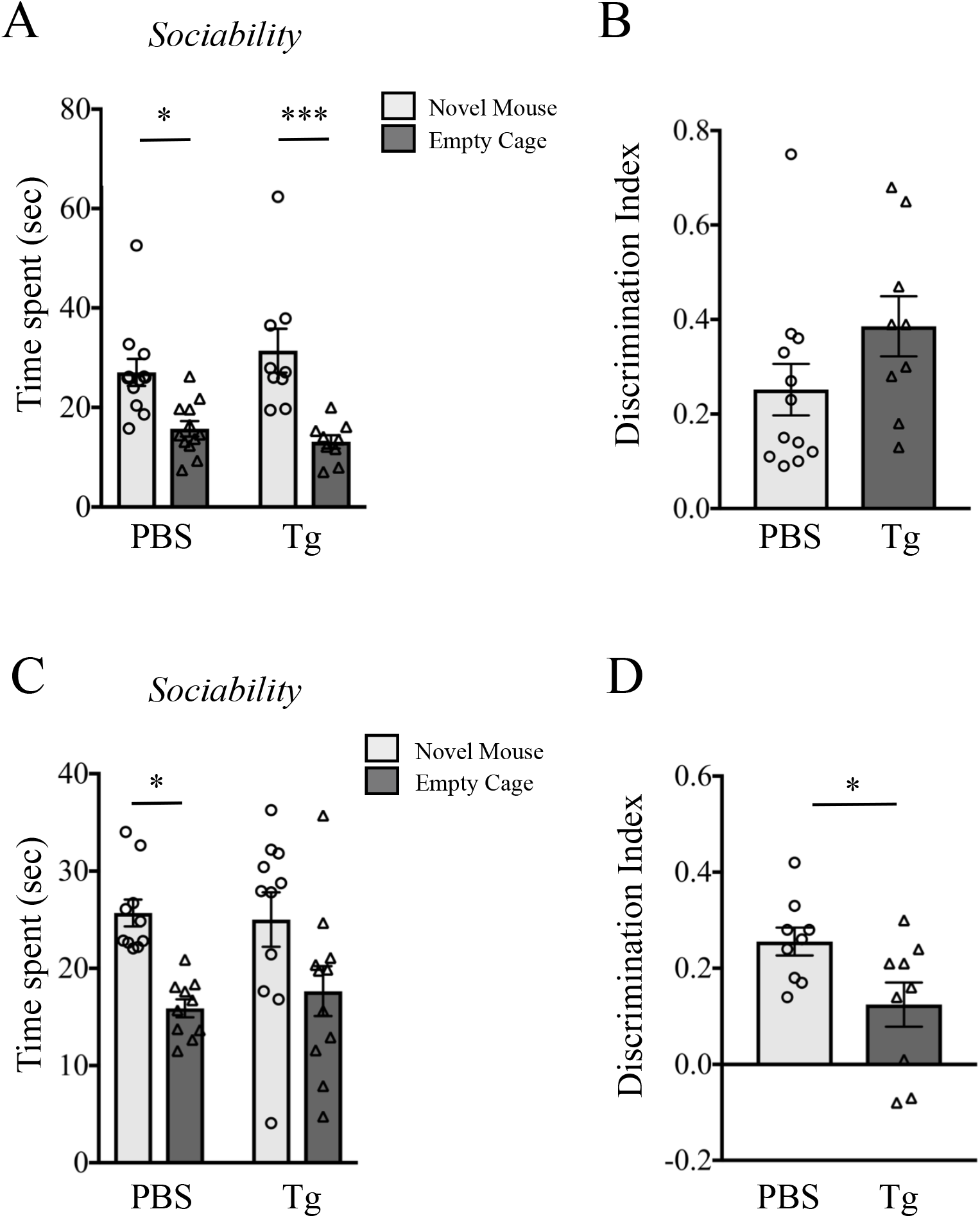
*T. gondii* infected mice display impaired sociability 8 weeks post infection (wpi), but not 3 wpi. Total time spent exploring an empty cage versus a cage containing a novel conspecific by *T. gondii* and mock infected mice was measured at 3 wpi (A) and 8 wpi (C). Discrimination index was calculated as the difference in time exploring the two cages divided by total exploration time at 3 wpi (B) and 8 wpi (D). In all graphs, error bars represent mean ± SEM (n= 9-12). Data were analysed by Student’s *t*-test (C and E) or twoway ANOVA with Sidak’s multiple comparison test as a *post hoc* (B and D). \**P*<0.05 and \*\*\**P*<0.001.

### 3.4 Infection impairs social recognition memory, but object recognition memory remains intact

Social recognition memory is a basic component of social behaviour and entails discrimination between a novel and familiar mouse (Berry and Bronson, 1992; Jiming et al., 1994; Kogan et al., 2000). A lack of social preference at 8 wpi (Fig.3 C,D) led us to hypothesise that at this stage of infection, mice would also display impaired social recognition memory. Moreover, brain regions involved in social memory, such as hippocampus and amygdala, are also involved in object recognition memory (Broadbent et al., 2010; Roozendaal et al., 2008). Therefore, using a separate cohort of mice, we investigated whether *T. gondii* infected mice displayed deficits in object recognition memory.

Using the 3-chambered social interaction chamber, we observed that, at 8 wpi, infected mice displayed impaired sociability by spending equal amount of time investigating an empty cage and a cage containing a mouse (mouse A), reinforcing our findings described above (Fig. 3). One hour later, when these mice were presented with a novel mouse (mouse B) in the presence of the familiar mouse (mouse A), the infected mice displayed no preference for the novel mouse (Fig. 4A), indicating impaired short-term social recognition memory. Moreover, the infected mice displayed significantly reduced discrimination index in the sociability test (*P* = 0.0011) and reduced recognition index in the memory recall phase (*P* = 0.0004), as compared to PBS mice (Fig. 4B). Interestingly, we noticed that *T. gondii* infection had no effect of short-term (1 hr) or long-term (24 hr) object recognition memory as they explored the novel object significantly more than the familiar one (*P* < 0.0001) (Fig. 4A), and displayed discrimination index comparable to PBS mice 1 hr as well as 24 hr after training (Fig. 4B). Thus, our results indicate that *T. gondii* infection alters a specific domain of learning and memory.

**Figure 4:**
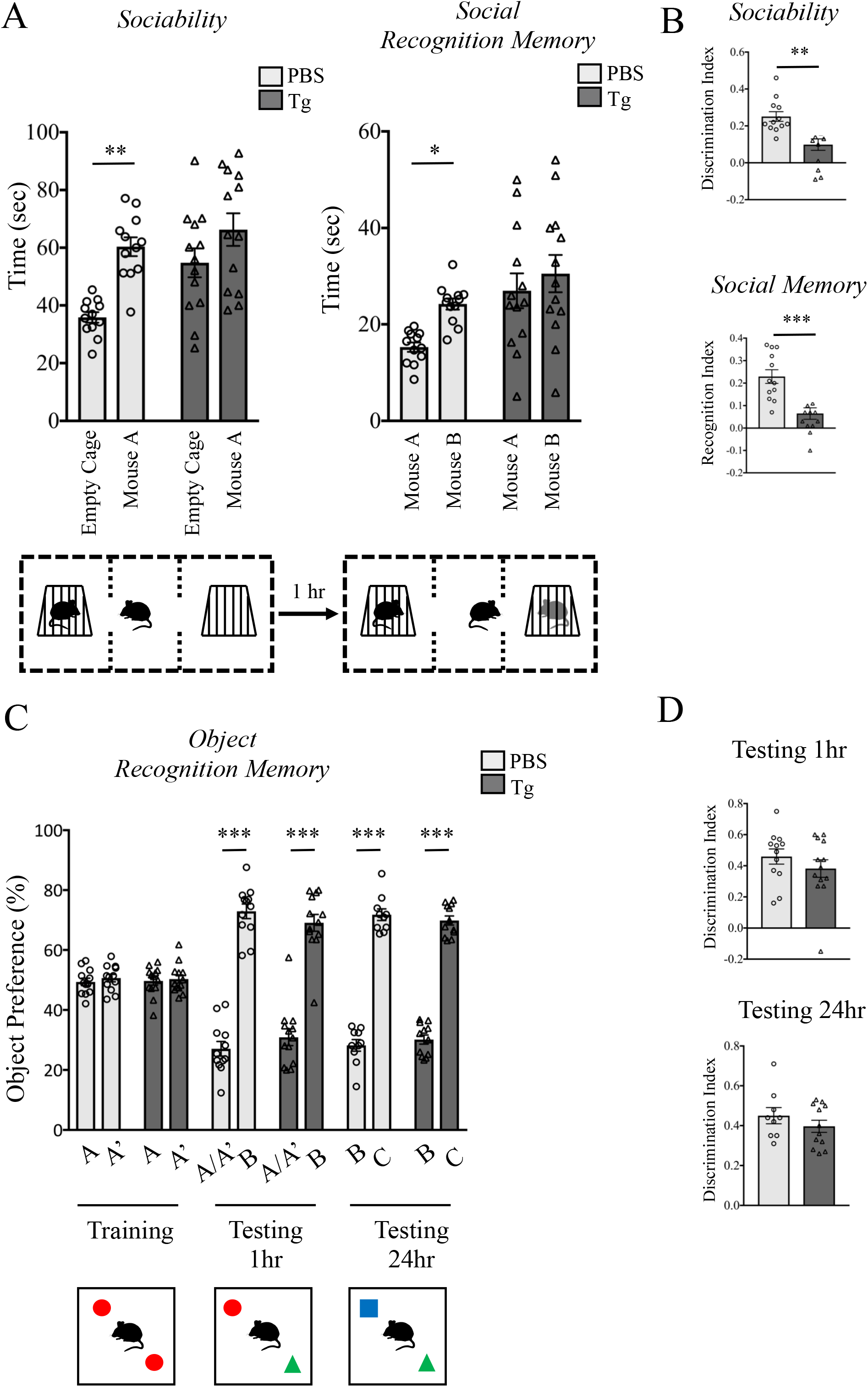
Chronic *T. gondii* infection impairs social, but not object recognition memory. (A) Sociability was measured as total time spent exploring an empty cage versus a cage containing a novel conspecific by *T. gondii* and mock infected mice at 8 wpi (A). To test the social recognition memory 1 hr later, total time spent exploring familiar versus novel mouse was measured. Discrimination indices for sociability and social memory were calculated as the difference in time spent exploring two cages divided by total exploration time (B). In the object recognition test, time spent exploring both objects was measured 1 hr (STM) or 24 hr (LTM) after training, and represented as a percentage of object preference (C). Discrimination indices were calculated as the difference in time exploring the two objects divided by total exploration time. In all graphs, error bars represent mean ± SEM (n= 12-13). Data were analysed by Student’s *t*-test (B and D) or two-way ANOVA with Sidak’s multiple comparison test as a *post hoc* (A and C). \**P*<0.05, \*\**P*<0.01 and \*\*\**P*<0.001.

### 3.5 *T. gondii* infection impairs neuronal activation in the brain

To identify changes in neuronal signalling that might be causing sociability deficits in mice 8wpi, we decided to map c-Fos induction in the brain regions that are known to be strongly involved in social behaviour. To generate c-Fos expression induced by social interaction, we exposed one group of infected and PBS mice to a novel mouse for 15 min (social) and left the other group in their home cage (basal). Two hours later, we performed immunohistochemistry on brain sections to measure c-Fos expression (Fig. 5A). We measured c-Fos positive cells in the hippocampus (CA1, CA2 and CA3 subfields), amygdala (basolateral (BLA) and central (CeA) regions) and the mPFC (prelimbic (PrL) and orbital cortex (OrbC) regions).

**Figure 5:**
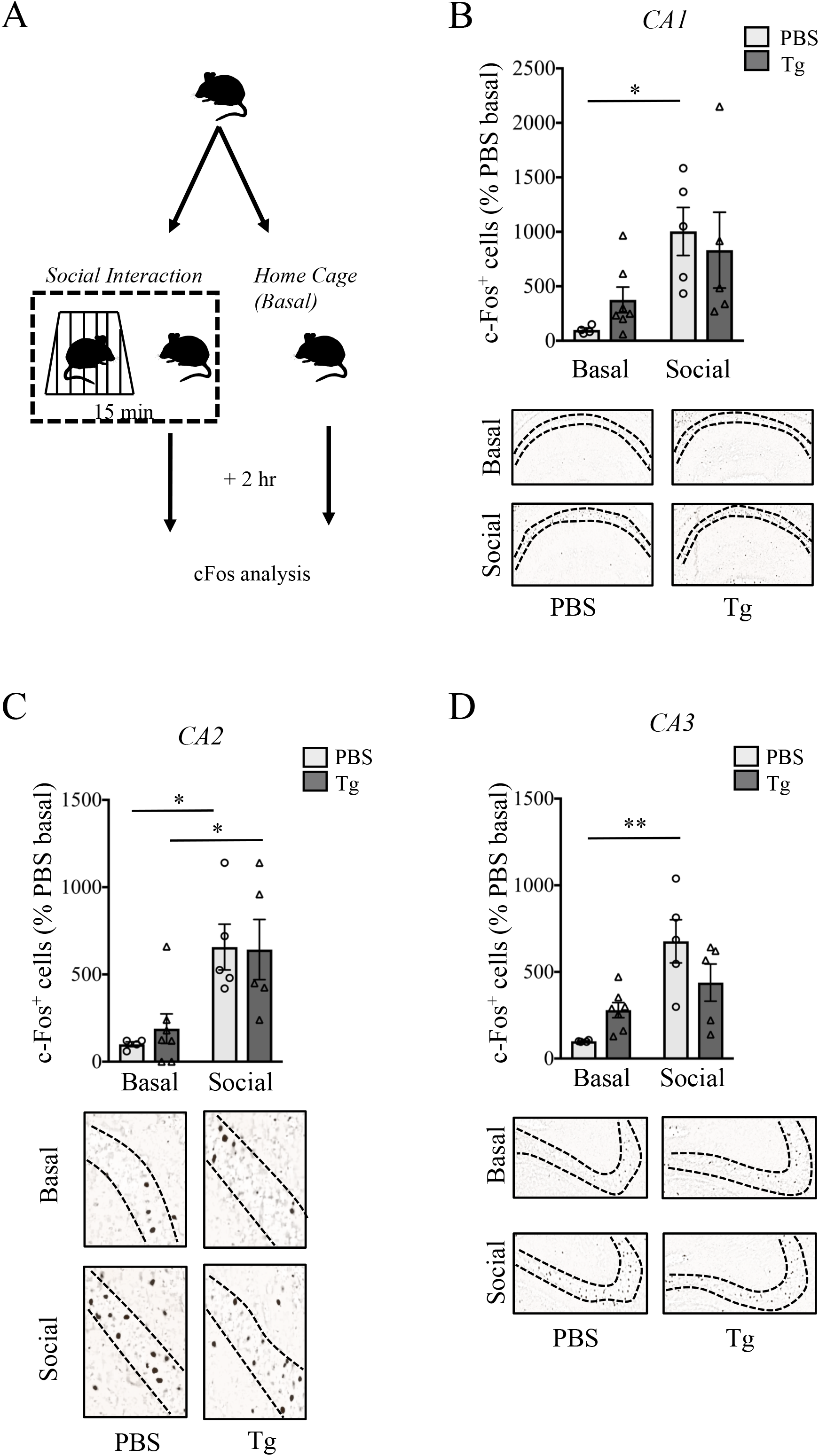
c-Fos expression in the hippocampus after exposure to a novel mouse. Schematic representation of the experimental setup to measure c-Fos expression in mice after social interaction (A). Number of c-Fos positive cells were measured in the CA1 (B), CA2 (C) and CA3 (D) regions of the hippocampus of PBS or *T. gondii* infected mice after social interaction or remaining in the home cage. c-Fos expression is displayed as a percentage of PBS mice remaining in the home cage. Representative images of immunohistochemical staining are shown. In all graphs, error bars represent mean ± SEM (n= 4-7). Data were analysed by two-way ANOVA with Sidak’s multiple comparison test as a *post hoc* (B-D). \**P*<0.05 and \*\**P*<0.01.

Two-way ANOVA revealed that PBS mice exposed to a social stimulus showed robust c-Fos induction in the CA1 (*P* = 0.0253), CA2 (*P* = 0.0138) and CA3 (*P* = 0.0019) regions of the hippocampus (Fig. 5B-D), the PrL (*P* = 0.0166) and OrbC (*P* = 0.0002) regions of the mPFC (Fig. 6A), and BLA (*P* = 0.0294) region of the amygdala (Fig. 6B), when compared to c-Fos positive cells in corresponding brain regions of PBS control mice that remained in the home cage. Although there appeared to be a small increase in the number of c-Fos positive cells in the CeA regions of the amygdala, the effect was not statistically significant (*P* = 0.6527). This indicates that these brain regions are involved in processing information when mice are exposed to novel social environment, as described previously. When we measured c-Fos positive cells in the hippocampus of infected mice, 2-way ANOVA revealed no significant c-Fos induction in the CA1 and CA3 regions when compared to c-Fos positive cells in corresponding brain regions of infected mice that remained in the home cage (Fig. 5B,D). We saw a similar effect in the OrbC region of the mPFC (Fig. 6A), and BLA and CeA regions of the amygdala (Fig. 6B). However, the CA2 (Fig. 5C) and PrL (Fig. 6A) regions showed significant increase in c-Fos expression in infected mice (*P* = 0.0214 and *P* = 0.0499, respectively) when exposed to a novel mouse. Interestingly, we observed that in the infected mice, the number of c-Fos positive cells in the CA1, CA3, and the BLA regions at basal levels were higher than that of PBS mice, although the effect of was not statistically significant.

**Figure 6:**
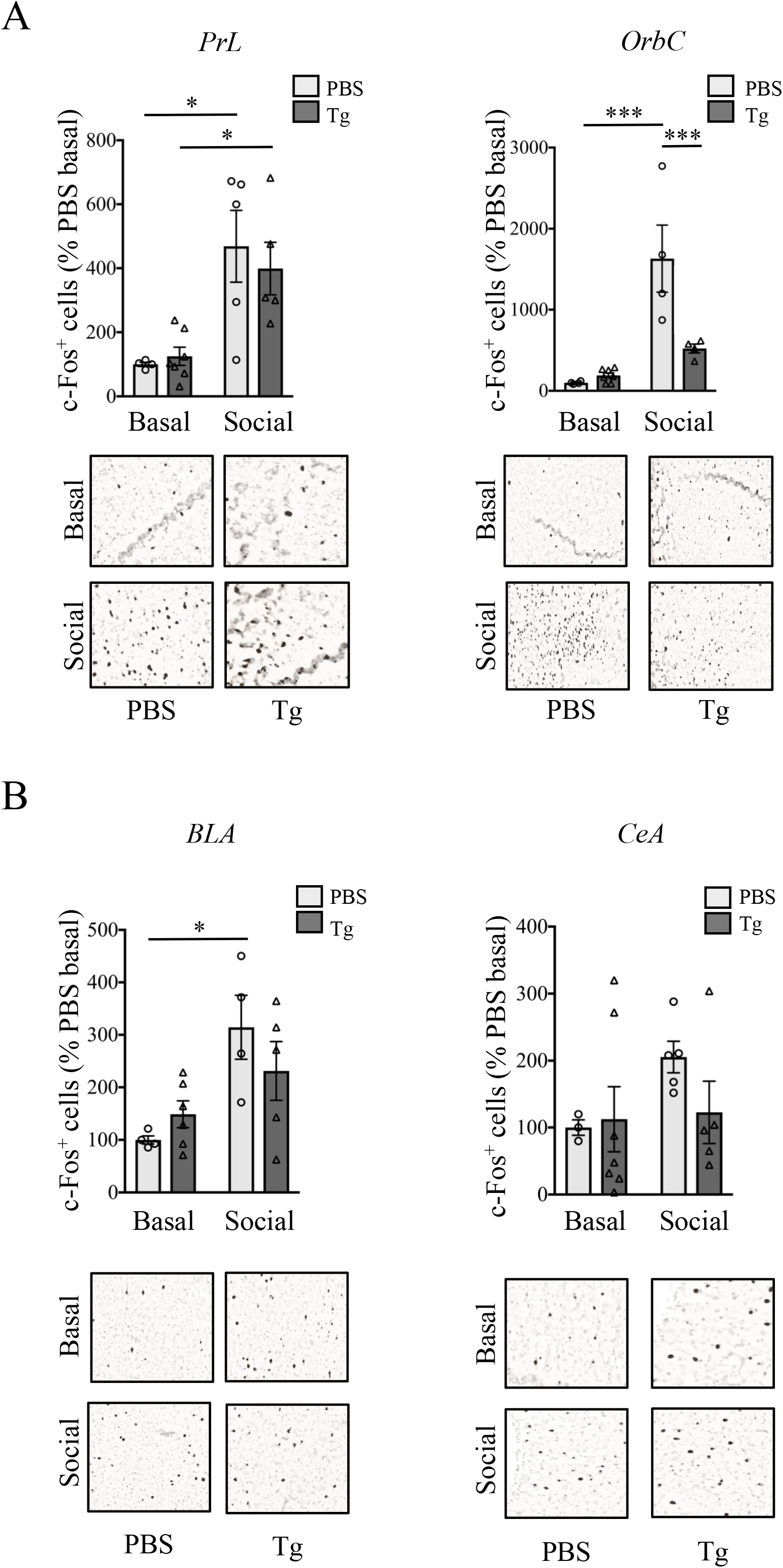
c-Fos expression in the mPFC and amygdala after exposure to a novel mouse. Number of c-Fos positive cells were measured in the PrL and OrbC regions of the mPFC (A) and CeA and BLA regions of amygdala (B) in PBS or *T. gondii* infected mice after social interaction or remaining in the home cage. c-Fos expression is displayed as a percentage of PBS mice remaining in the home cage. Representative images of immunohistochemical staining are shown. In all graphs, error bars represent mean ± SEM (n= 4-7). Data were analysed by two-way ANOVA with Sidak’s multiple comparison test as a *post hoc* (B-D). \**P*<0.05, \*\**P*<0.01 and \*\*\**P*<0.001.

Our results indicate that: (1) *T. gondii* infection impairs IEG expression in the hippocampus, mPFC and amygdala when mice are exposed to social stimuli but the effect is heterogeneous, in that, some regions are spared and do not show any impairment in IEG induction, (2) impaired social recognition memory could be due to impaired neuronal activation upon social interaction, and (3) changes in the way neurons in these brain regions react to incoming stimuli does not affect object recognition memory.

### 3.6 Functional connectivity between brain regions after social interaction

Out results mapping c-Fos activity upon social interaction revealed changes in neural activity in multiple brain regions. Indeed, memory is stored in a network comprising of multiple brain regions (Fukushima et al., 2014; Tanimizu et al., 2017; Wheeler et al., 2013; Zhang et al., 2011) and an understanding of changes in circuits that modulate social behaviour can provide deeper insights into behavioural changes upon *T. gondii* infection. We calculated covariance within brain regions using the results of c-Fos expression from Figure 5 and 6 to infer strength of interaction between neural circuits (Horwitz et al., 1995; McIntosh, 1999). Figure 7 shows Pearson’s *r* correlation matrices for PBS and *T. gondii* infected mice at basal (home cage) level or 2 hr post social interaction.

**Figure 7:**
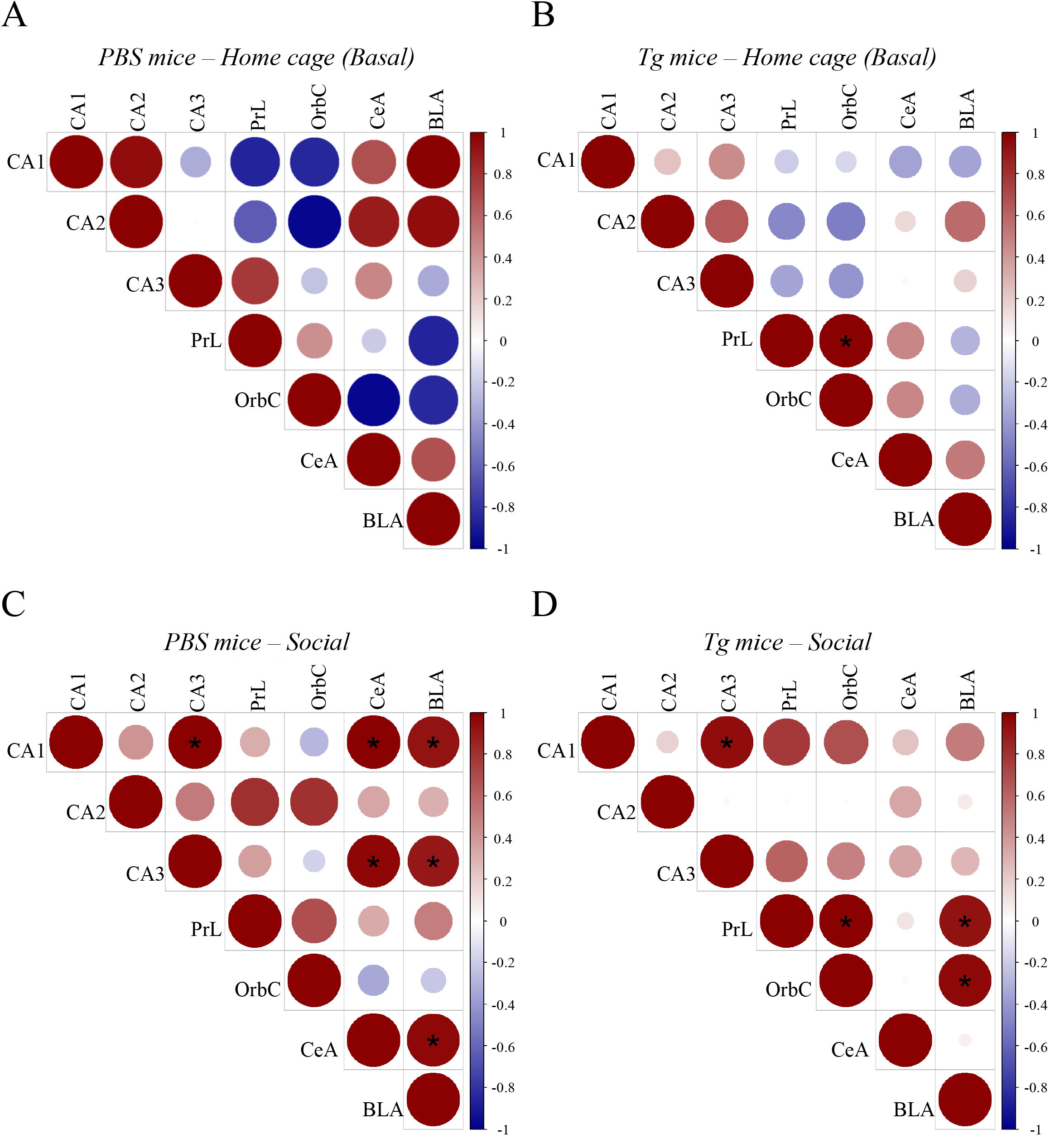
Functional connectivity between brain regions after social interaction. Matrices show interregional correlations for c-Fos expression in PBS mice in their home cage (A) or after social interaction (C) and *T. gondii* infected mice in their home cage (B) or after social interaction (D). Colour represents correlation strength based on Pearson’s *r* values (scale shown on the right), while the size of the circle represents absolute value of the correlation coefficient. Data were analysed using the R Hmisc package. \**P*<0.05 compared within the group.

Correlation analysis showed that pattern of interconnectivity between brain regions is altered in the home cage *T. gondii* infected mice, as compared to PBS mice (Fig. 7A). When exposed to a novel conspecific, Pearson’s *r* value for c-Fos expression among all brain regions of the PBS mice showed an increased positive correlation, with a significant correlation (P < 0.05) between the hippocampus and the amygdala as well as within the subregions of the hippocampus and amygdala (Fig. 7B), suggesting a strong connectivity in these regions when mice interact in a social environment. In the *T. gondii* infected mice, although we observed an overall increase in positive correlations among brain region after social interaction, such correlations are different from the PBS mice. There was significant positive correlation (*P* < 0.05) between the mPFC and the amygdala in the *T. gondii* mice and within the mPFC, which was absent in the PBS mice (Fig. 7B).

Our results show that functional connectively between brain regions is altered upon *T. gondii* infection, which could change how information is processed and stored when social recognition memory is generated.

### 3.7 Changes in the components of neuronal signalling in the hippocampus, amygdala and PFC of *T. gondii* infected mice

We investigated the molecular changes underlying the observed behavioural changes in chronic *T. gondii* infected mice. The role of cyclic-AMP/PKA signalling at the synapse in learning and memory formation is well established (Abel et al., 1997; Abel and Nguyen, 2008; Arnsten et al., 2005), and in this process, several other proteins play a role in maintaining the synaptic integrity as well as in mediating the downstream signalling cascade involved in gene transcription implicated in long-term memory formation. External stimuli, such as learning or social interaction, increases neuronal activity that induces c-Fos (Flavell and Greenberg, 2008). Interestingly, activity regulated intracellular [Ca^2+^], that mediates c-Fos gene expression (Ghosh et al., 1994; Thompson et al., 1995), also can activate Ca^2+/^ calmodulin dependent kinases (CAMK) that regulate adenylyl cyclase activity and cAMP-PKA-CREB signalling (Halls and Cooper, 2011; Poser and Storm, 2001). Because we found impaired c-Fos induction upon learning, we investigated whether the basal cAMP pools in these brain regions were also affected upon infection. Immunofluorescence staining of cAMP revealed that in *T. gondii* infected mice, total cAMP was reduced in the CA1 (*P* = 0.003) and CA3 (*P* = 0.0296) regions of the hippocampus, but not the CA2 region (Fig. 8A). In contrast, cAMP levels were significantly elevated in the amygdala (*P* = 0.0003) and the PrL (*P* = 0.03) of mPFC (Fig. 8B,C), but remained unchanged in the OrbC (Fig. 8B), indicating a region-specific regulation of cAMP levels due to *T. gondii* infection in the brain.

**Figure 8:**
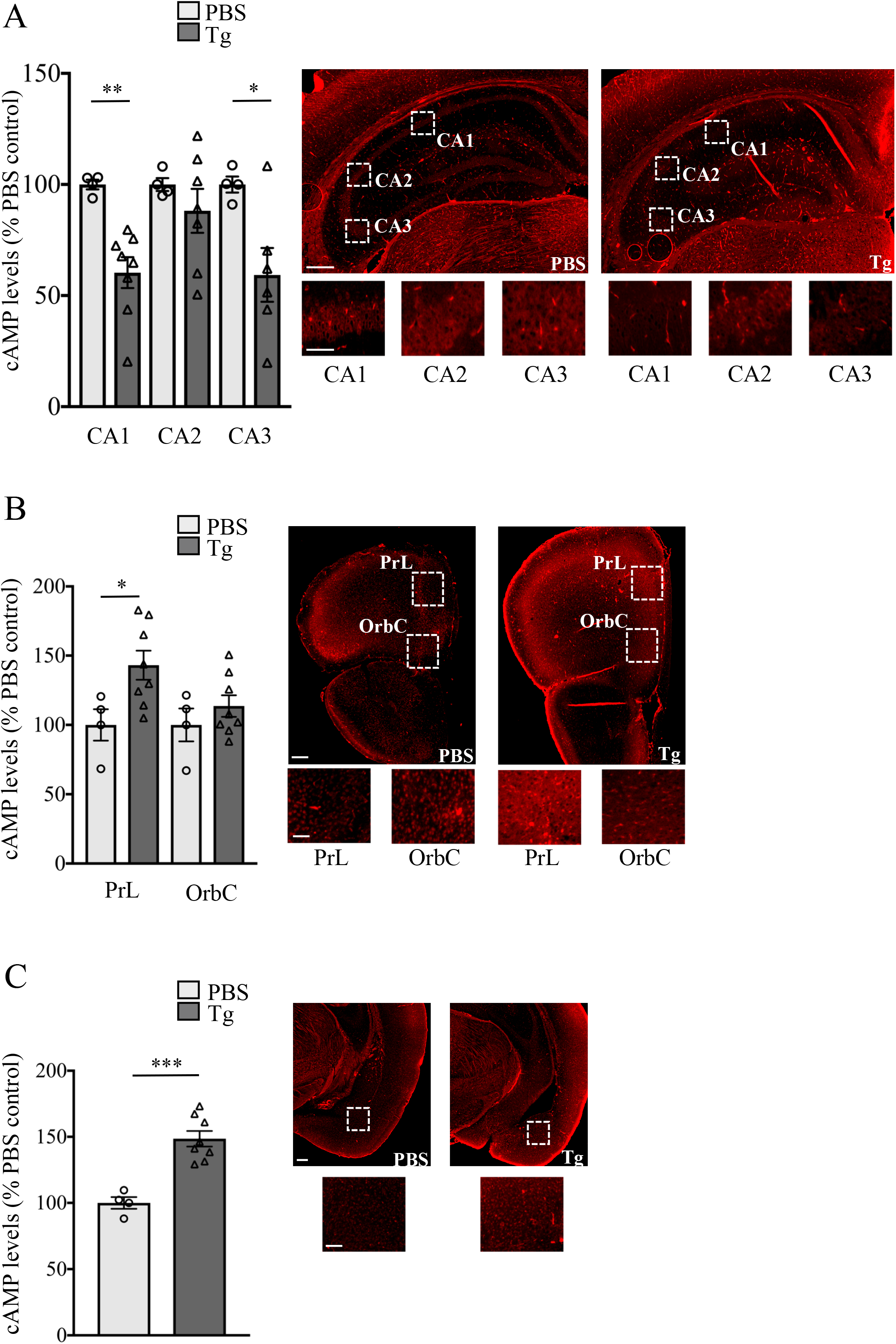
Chronic *T. gondii* infection alters brain cAMP levels. cAMP was analysed by immunohistochemistry of brain slices obtained from PBS injected or *T. gondii* infected mice 8 wpi. Graphs show the quantification of the average intensity of cAMP immunoreactivity from the CA1, CA2 and CA3 regions of the hippocampus (A), PrL and OrbC regions of the mPFC (B) and the CeA and BLA regions of amygdala (C). Values are represented as percentage of PBS mice. Representative photomicrographs with high magnification inset are shown for reach brain region. Scale bar 200 and 40 μm for low and high magnification images, respectively. Data were analysed by Student’s *t*-test. \**P*<0.05, \*\**P*<0.01 and \*\*\**P*<0.001.

Next, we measured levels of pre- and post-synaptic proteins that are involved in maintaining synaptic structural and functional integrity. Synaptophysin, a pre-synaptic protein involved in the regulation of vesicle fusion and recycling (Valtorta et al., 2004), was reduced (*P* = 0.0326) in the hippocampus of *T. gondii* infected mice (Fig. 9A). PSD95, that mediates correct assembly of postsynaptic density complex (Chen et al., 2011; Montgomery and Madison, 2004) and regulates synaptic plasticity, was significantly reduced in the hippocampus (*P* = 0.023) and PFC (*P* < 0.0001) (Fig. 9A,B).

**Figure 9:**
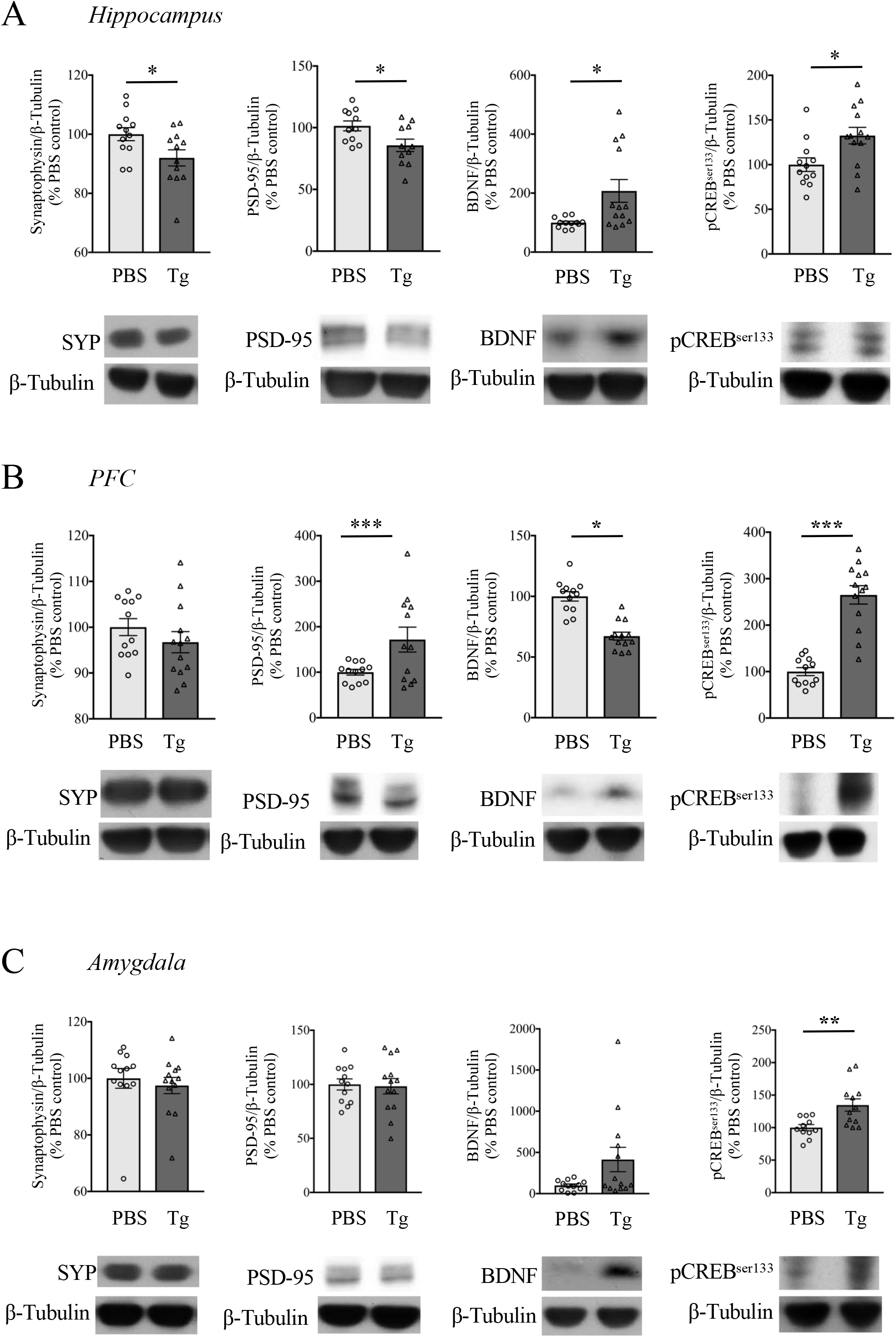
Changes in neuronal proteins after a chronic *T. gondii* infection. Synaptophysin, PSD-95, BDNF and phospho-CREB^ser133^ (pCREB^ser133^) levels were analysed by western blots of protein extracts obtained from the hippocampus (A), PFC (B) and amygdala (C) of PBS and *T. gondii* infected mice 8 wpi. Representative immunoblots are shown. Values obtained by densitometric analyses of western blot data are expressed as percentage of PBS-injected mice. In all graphs, error bars represent mean ± SEM (n= 11-12). Data were analysed by Student’s *t*-test. \**P*<0.05, \*\**P*<0.01 and \*\*\**P*<0.001.

We also measured proteins downstream of the cAMP-PKA signalling, including BDNF and phospho-CREB. BDNF, a neurotrophin secreted in response to activity (Matsuda et al., 2009) and involved in a wide range of neural function such as survival, development and plasticity (Binder and Scharfman, 2004; Huang and Reichardt, 2001; McAllister et al., 1999), was increased in the hippocampus (*P* = 0.0124) and PFC (*P* = 0.0184) (Fig. 9A,B). Interestingly, we found that levels of phosho-CREB, a regulator of activity-dependent gene transcription (Bourtchuladze et al., 1994; Josselyn et al., 2004; Kida et al., 2002; Pittenger et al., 2002), was increased in hippocampus (*P* = 0.0133), amygdala (*P* = 0.0047) as well as the PFC (*P* < 0.0001) (Fig. 9A-C).

Cumulatively, our results indicate that *T. gondii* infection leads to molecular changes in the hippocampus, amygdala and the prefrontal cortex which could fundamentally alter the way these brain regions acquire and consolidate external stimuli, leading to the chronic learning impairments we observed (8 wpi).

## 4. Discussion

In the present study, we report early changes in mice infected with *T. gondii* akin to sickness-induced behavioural alterations, while at chronic stages of infection, mice also begin to display cognitive impairments. We further demonstrate that at chronic stages, *T. gondii* infection leads to reduced sociability in mice and an impaired short-term social recognition memory but object recognition memory remained unaltered. This could be a result of impaired neuronal activation and altered functional connectivity among brain regions strongly associated with social behaviours in rodents (hippocampus, amygdala and mPFC) when placed in a social environment. We also demonstrate changes in molecules that play important role in synaptic plasticity and downstream signalling involved in learning and memory.

This is the first study to perform a detailed analysis of behavioural changes in *T. gondii* infected mice as early as 3 wpi. *T. gondii* cysts can be found in the brain at 3 wpi (Watts et al., 2015) and tachyzoites even earlier (Vyas et al., 2007). We found characteristic sickness-induced behaviour in mice such as increased anxiety, lethargy, anhedonia and depression-like phenotype which could be due to upregulation of pro-inflammatory molecules such as tumour necrosis factor-alpha (TFN-a), interleukin-12, and interleukin1B (Scanga et al., 2002) as well as activation of tryptophan/kynurenine metabolism pathway involving gamma interferon (IFN*γ*), indoleamine 2,3-dioxygenase, quinolinic acid and inducible nitric oxide synthase (Fujigaki et al., 2003, 2002; Mahmoud et al., 2017) all of which are known to induce behavioural changes that we observed. But, we failed to observe any changes in spatial memory and working memory of infected mice (at 3 wpi) when tested in a Y-maze. This is interesting because sickness behaviour induced by lipopolysaccharide (LPS) does lead to impairments in working memory (results not shown; also Parrott et al., 2016; Sparkman et al., 2006), and toll-like receptor 4 (TLR4), the major receptor for LPS, is involved in *T. gondii* recognition by the host (Zare-Bidaki et al., 2014). However, *T. gondii* also upregulates production of anti-inflammatory IL-10 (Khan et al., 1995; Neyer et al., 1997; Wilson et al., 2005) that might counteract TLR4 signalling effects on behaviour in our models. Here, we observe that at chronic stage of infection, mice display impaired spatial memory in the Y-maze, but their working memory remains intact. Previous studies provide conflicting results on Y-maze performance, which suggests that the effect might be strain dependent (Hutchison et al., 1980; Jung et al., 2012; Kannan et al., 2010; Xiao et al., 2016). Our results are in line with previous reports showing no working memory deficit, while providing first evidence of impaired spatial memory in the Y-maze caused by infection with the Pru strain of parasite. Nevertheless, because generalised inflammatory response due to pathogen infection, which is known to persist at chronic stages (Aliberti, 2005; Yarovinsky, 2014), is involved in such behavioural changes, it might be difficult to determine whether these changes are caused by direct parasitic activity.

Clinical and pre-clinical data has revealed that general anxiety disorders often exhibit comorbidity with social dysfunction (Allsop et al., 2014). Autistic patients have higher levels of reported anxiety (White et al., 2009), while people with anxiety disorders score higher on autism spectrum disorder symptoms scale (Pine et al., 2008). Also, drugs prescribed for generalised anxiety have also been used to treat social anxiety (Jefferson, 2001). Mouse models for autism display pronounced social as well as anxiety disorders (Nakatani et al., 2009; Peça et al., 2011; Silverman et al., 2010). Therefore, we hypothesised that the anxiogenic effect of *T. gondii* infection observed by us and others (Gatkowska et al., 2012; Hay et al., 1984; Mahmoudvand et al., 2015; Skallová et al., 2006; Torres et al., 2018) could lead to changes in social behaviours in infected mice. Previously, Gonzalez et al. showed that lower doses of *T. gondii* infection (100 and 1000 tachyzoites) impaired social interaction in rats, but the effect was lost at higher doses. Only one recent study (Torres et al., 2018) has investigated social behaviour in mice infected with *T. gondii*, but report no changes in social interaction, despite increased anxiety. We evaluated sociability in mice using the 3-chamber sociability test at 3wpi, as well as 8pwi, and observed an impaired sociability only at 8wpi. Discrepancy between our results and Torres et al. could be due to the parasite strain or their route/dose of infection (10 ME49 cysts, orally). However, despite increased anxiety at 3 wpi, we do not observe any impairment in social interaction. This result points towards our previous assumption that behavioural changes at the acute stage could be a result of general inflammatory response which might not include social dysfunction, and propound the idea that at a chronic stage, this effect could be due to long-term changes in neuronal function either caused by parasite itself, or a long-term effect of initial immune response in the brain, or both. It is interesting to note that in the acute infection stage, there is a strong upregulation of IFN*γ* in the brains of infected mice (Mahmoud et al., 2017) which is critical for host survival (Sa et al., 2015; Sturge and Yarovinsky, 2014) and IFN*γ* plays important reinforcing role in sociability in mice (Filiano et al., 2016) which could explain our results at 3 wpi. Nevertheless, it seems that a chronic *T. gondii* infection which is accompanied by increased anxiety can lead to impaired sociability.

When we measured social recognition memory in *T. gondii* infected mice, we observed impaired social recognition STM. Recently, Torres et al. were the first ones to show social recognition impairment in *T. gondii* infected mice (Torres et al., 2018). That study presented combined data from female and male mice and therefore results could be confounded due to reported sex differences in *T. gondii* infection, as shown previously (Gatkowska et al., 2013; Roberts et al., 1995; Xiao et al., 2012). Interestingly, we observed that in these mice, object recognition STM as well as LTM was intact. Previously, Gulinello et al. has also reported intact object recognition (Gulinello et al., 2010), while later, Xiao et al. reported that only the mice displaying high MAG1 antibody levels show impairment in long-term object recognition memory (Xiao et al., 2016). Different parasite strain and mouse genotypes used in these studies could account for such inconsistencies. However, we also report an impaired spatial recognition memory in the Y-maze in this study. This suggests differential influence of *T. gondii* infection on performance, which could arise due to the heterogeneity in the function of the structures of the temporal lobe as described by previous studies (Balderas et al., 2008; Forwood et al., 2005; Winters et al., 2004). Further experiments are required to determine if impaired neuronal activation is the cause or effect of social dysfunction in *T. gondii* infected mice. Furthermore, intact object recognition memory indicates that although there is an overlap of brain regions involved in these two types of recognition memories, *T. gondii* infection leads to a preferential social impairment.

Rodent social behaviour is a complex trait involving a crosstalk between multiple brain regions and various signalling pathways that regulate behaviours such as social approach, social novelty and ability to discriminate familiar and novel conspecifics. Monitoring activation of brain regions involved in these behaviours can give clues to the signalling dysfunction caused by *T. gondii*. Moreover, non-invasive imaging has revealed that *T. gondii* infection causes significant changes in neuronal pathology, leading to a loss of grey matter, reduced fiber coherence and lesions in the brain (Horacek et al., 2012; Parlog et al., 2014). Therefore, we quantified induction of immediate early gene c-Fos in the hippocampus, amygdala and the mPFC when mice were presented with a social stimulus and used that data to compute functional correlations between brain regions. At basal levels, low c-Fos levels are maintained due to instability of c-Fos mRNA and auto-repression of c-Fos transcription by the Fos protein itself (Lucibello et al., 1989; Morgan and Curran, 1991). Therefore, it is suggested that a short-term sensory deprivation in the form of social isolation prior to stimulus presentation leads to the best activity dependent c-Fos signal (Chung, 2015). Interestingly, several studies have highlighted that long-term socially isolated rodents fail to show a robust social interaction induced c-Fos production (Ieraci et al., 2016; Lukkes et al., 2012; Wall, 2012). In contrast, it has also been shown that isolation-housed rats approached a novel conspecific more than group-housed rats, and isolation acted as a strong motivator to social approach (Templer et al., 2018; Wall, 2012). Indeed, a 3 week isolation period did not impair c-Fos induction upon social intersection (Avale et al., 2011). Accordingly, in this study, we single housed mice 3-4 days prior to c-Fos induction experiment such that it motivated social interaction but did not impair c-Fos activation, as it did after a chronic social isolation. Here, we observe a robust c-Fos activation in the hippocampus (CA1, CA2 and CA3), mPFC (PrL and OrbC) and amygdala (BLA and CeA) of PBS mice when presented with a social stimulus. However, *T. gondii* infected mice displayed an impaired c-Fos induction in all regions, except the CA2 and PrL. Also, we observed changes in pattern of connectivity between brain regions between PBS controls and *T. gondii* mice at rest in their home cage. Our results contradict previous studies that report no IEG induction in the CA2 region of the hippocampus upon social stimuli (Alexander et al., 2016; Tanimizu et al., 2017), while supports findings from Kim et al. that report CA2 activation upon social interaction (Kim et al., 2015). The hippocampus is critically integrated into the rodent circuitry involved in social behaviours (Kim et al., 2015; Ko, 2017; Kogan et al., 2000; Raam et al., 2017; Rubin et al., 2014), but specifically, the CA2 subfield is important for the formation of social memory (Alexander et al., 2016; Hitti and Siegelbaum, 2014; Leroy et al., 2017; Lin et al., 2017; Smith et al., 2016; Stevenson and Caldwell, 2014). Therefore, it is no surprise that we observe neuronal activation in the CA2 region upon social stimuli, a prerequisite to social memory formation (Felix-Ortiz and Tye, 2014; Hitti and Siegelbaum, 2014). However, intact c-Fos induction in CA2 region of infected mice suggests that *T. gondii* infection affects a specific set of neurons and corroborates with the distinct anatomical and physiological make up of neurons from the CA1/CA3 and CA2 regions (Dudek et al., 2016; Ko, 2017). Furthermore, our results indicate that a lack of social memory in *T. gondii* mice could result from an altered connectivity of CA2 region with the mPFC and amygdala. Social interaction also involves activation of mPFC (Jodo et al., 2010; Yizhar et al., 2011) and the amygdala (Katayama et al., 2009; López-Cruz et al., 2017) and lack of a robust c-Fos induction in these regions in *T. gondii* infected mice, along with altered functional correlation among brain regions, suggests impairments in social-behaviour related neural circuits in the mice brain. Other factors that could affect c-Fos induction in infected animals is presence of a chronic anxiety phenotype (File and Seth, 2003) and a sustained elevation of anti-NMDAR antibodies (Kannan et al., 2017, 2016; Lucchese, 2017) or possibly loss of NMDARs (Torres et al., 2018) that could impair ion-channel gated c-Fos induction during neuronal activity (Herdegen and Leah, 1998). Nevertheless, more detailed studies are required to differentiate the specific neural subtypes or pathways that undergo preferential dysfunction upon *T. gondii* infection.

During synaptic activity, NMDARs and L-type VSCC mediate the increase in intracellular [Ca^2+^] concentration that engages c-Fos transcription (Chaudhuri et al., 2000; Ghosh et al., 1994), mainly via the MAPK pathway (Chaudhuri et al., 2000). Concurrently, this also activates CAM kinases and CAM-stimulated adenylyl cyclase 1 and 8 (Xia and Storm, 2012), that regulate the cAMP pools and therefore PKA signalling, all of which are critical in learning and memory formation. Ability of *T. gondii* to modulate MAPK (Braun et al., 2013) and NMDAR signalling (Kannan et al., 2017, 2016; Torres et al., 2018) are previously described. MAPK signalling phosphorylates CREB at ser133 (Impey et al., 1998) that regulates c-Fos (Sheng et al., 1990) as well as BDNF (Tao et al., 1998) transcription. In the hippocampus of *T. gondii* infected mice, we found significant reduction in cAMP pools in the CA1 and CA3, but not in the CA2 subfield, which could be due to altered synaptic integrity and functionality, indicated by deceased PSD95 and synaptophysin levels in our results and elsewhere (Parlog et al., 2014). Our current findings are consistent with known functional and structural differences in the CA1/CA3 and CA2 regions (Dudek et al., 2016). CA2 neurons are more involved with object recognition (Lee et al., 2010), and carry less spatial information compared to CA2/CA3 neurons (Alexander et al., 2016; Mankin et al., 2015); consistent with intact object recognition and impaired spatial memory in our mice. Moreover, we find upregulated basal cAMP levels in the amygdala and mPFC. Interestingly, long-term increases in cAMP signalling impair social recognition memory (Garelick et al., 2009) and lead to cognitive impairments (Tyebji et al., 2015). Increased cAMP levels could also indicate a possible protein kinase A mediated anti-inflammatory mechanism (Brown et al., 2011; Power Coombs et al., 2011) to counteract toll-like receptor mediated inflammation in response to *T. gondii* (Debierre-Grockiego et al., 2007; Yarovinsky, 2014; Zare-Bidaki et al., 2014). We find that BDNF levels in the hippocampus and the PFC are increased after infection. Previous studies have shown that levels of BDNF, as well its regulator MiR132, are upregulated in acute *T. gondii* infection (Xiao et al., 2014; YongHua et al., 2009) but during chronic infection, only BDNF levels were found to be increased while MiR132 levels normalise (Li et al., 2015).

In the present study, we conclude that social behaviour in mice during chronic *T. gondii* infection is impaired which could arise due to dysfunctional neuronal signalling in regions of the brain relevant for social behaviour in rodents. Our results indicate that *T. gondii* infection affects a specific set of behavioural domains while others are spared, and therefore it is tempting to speculate that the effects of *T. gondii* infection on the brain could be specific to some neuronal subtypes/pathways. Further experiments are required to elucidate brain region/cell type specific functional changes, and thus find molecular targets to reverse the cognitive impairments observed during the chronic infection.

## Supporting information

## Author contributions

ST, SS, CJT, Conception and experimental design, Data acquisition, Interpretation and analysis of data, Drafting and revising the article; AJH, Conception and experimental design, Interpretation and analysis of data, Drafting and revising the article; ALG, Interpretation and analysis of data, Drafting and revising the article.

## Acknowledgements

We would like to thank the expert help of Carolina Alvarado from WEHI Bioservices, Onker Singh from WEHI Engineering department, Ellen Tsui and Cary Tsui form WEHI Histology department, and Lachlan Whitehead and Mark Scott from WEHI’s Centre of Dynamic Imaging. We would also like to thank Emma Burrows and Shlomo Yeshurun from Florey Institute of Neuroscience and Mental Health, Melbourne for their useful discussions and support for this work. SS is a recipient of the Australian Government Research Training Program Stipend Scholarship. AJH is an NHMRC Principal Research Fellow. We also gratefully acknowledge The David Winston Turner Endowment for funding this work. We are also grateful for institutional support from the Victorian State Government Operational Infrastructure Support and the Australian Government NHMRC IRIISS.

