## Supplementary material for "Impaired social behaviour and molecular mediators of associated neural circuits during chronic *Toxoplasma gondii* infection in female mice"

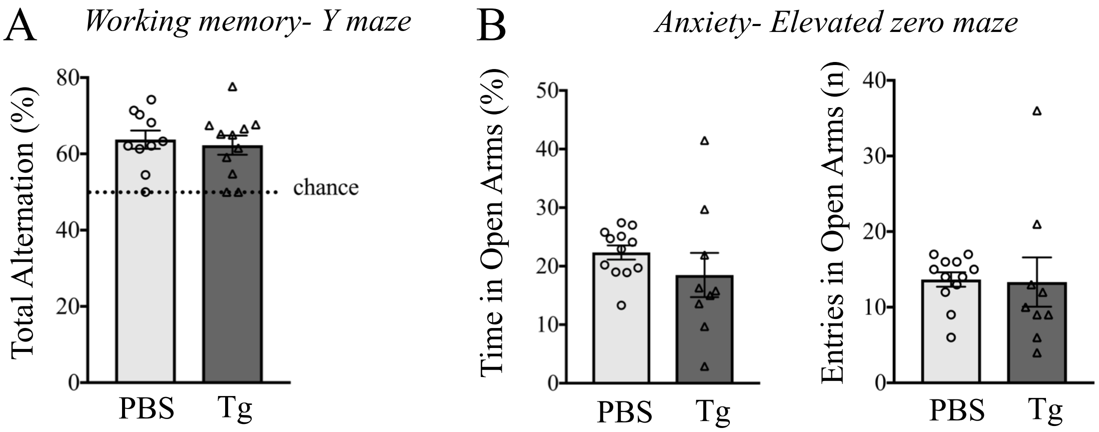


**Supplementary Figure 1: Changes in anxiety and working memory in mice 3 weeks after *T. gondii* infection.** 6 to 8-week-old female C57BL/6 mice were infected with *T. gondii* (Tg) or mock infected (PBS) and tested 3 weeks later. Working memory was tested by calculating the percentage alternation on the Y-maze continuous alternation test (A). Anxiety was measured using the elevated zero maze by measuring the time spent and entries made into the open zones (B). In all graphs, error bars represent mean ± SEM (n= 9-12). Data were analysed by Student's *t*-test.

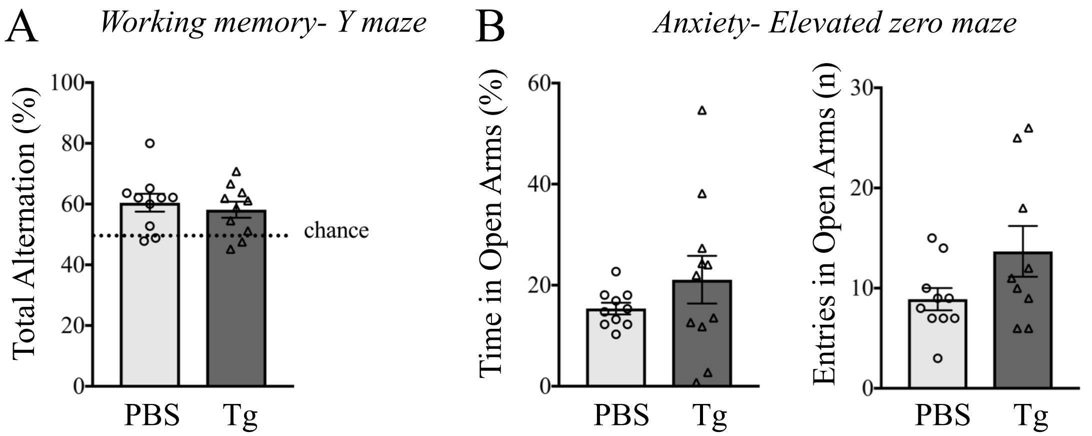


**Supplementary Figure 2: Changes in anxiety and working memory in mice 8 weeks after *T. gondii* infection.** 6 to 8-week-old female C57BL/6 mice were infected with *T. gondii* (Tg) or mock infected (PBS) and tested 8 weeks later. Working memory was tested by calculating the percentage alternation on the Y-maze continuous alternation test (A). Anxiety was measured using the elevated zero maze by measuring the time spent and entries made into the open zones (B). In all graphs, error bars represent mean ± SEM (n= 8-11). Data were analysed by Student's *t*-test.

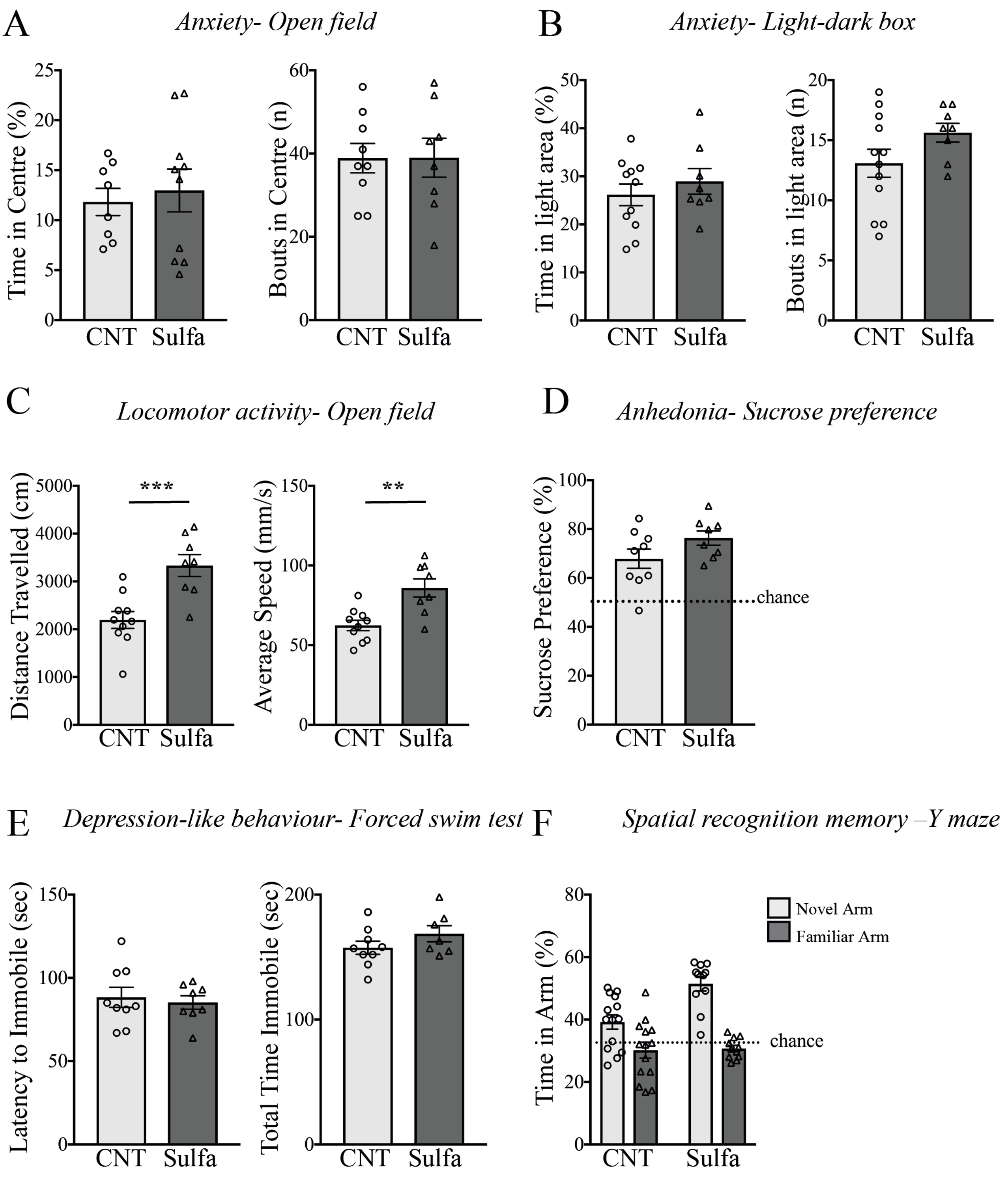


**Supplementary Figure 3: Effect of sulfadiazine treatment on mouse behavioural.** 6 to 8-week-old female C57BL/6 mice were provided water (CNT) or sulfadiazine (Sulfa) in drinking water (100 μg/ml) for 5 days and behaviour analysis began 2 weeks later. Anxiety phenotype was measured using the open field test (A) and the light-dark box test (B) by calculating the percentage times spent and entries made into the centre area or the light-zone, respectively. Locomotor activity was measured as the distance travelled and average speed in the open field (C). Anhedonia was measured by calculating preferential sucrose consumption over water (D). Depression-like behaviour was analysed by measuring the latency to immobility and the total time immobile in the forced-swim test (E). Spatial recognition memory was measured using the Y-maze as time spent in each arm 1 hour after training (F). In all graphs, error bars represent mean ± SEM (n= 8-11). Data were analysed by Student's *t*-test (A-E) or two-way ANOVA with Sidak's multiple comparison test as a *post hoc* (F). $^{**}P<0.01$ and $^{***}P<0.001$.
